# Repurposing the dark genome. II - Reverse Proteins

**DOI:** 10.1101/2023.03.20.533367

**Authors:** Sarangadhar Nayak, Pawan K. Dhar

**Author notes:** Corresponding author Pawan K. Dhar, Ph.D., Professor & Head, Synthetic Biology group, School of Biotechnology, Jawaharlal Nehru University New Delhi 110067, web: https://www.syntheticbiology.in.

## Abstract

Based on the expression blueprint encoded in the genome, three groups of sequences have been identified – protein encoding, RNA encoding, and non-expressing. We asked: Why did nature choose a particular DNA sequence for expression? Did she sample every possibility, approving some for RNA synthesis, some for protein synthesis, and retiring/ignoring the rest. If evolution randomly selected sequences for metabolic trials, how much non-utilized (not-expressing) and under-utilized (only RNA encoding) information is currently available for innovations? These questions lead us to experimentally synthesizing functional proteins from intergenic sequences of E.coli (Dhar et al 2009). The current work is an extension of this original report and takes into consideration natural protein-coding sequences ‘read backward’ to generate a new possibility. Reverse proteins are full-length ‘translation equivalents’ of the existing protein-coding genes read in the -1 frame. The structural, functional and interaction predictions of reverse proteins in *E*.*coli, S*.*cerevisiae* and *D*.*melanogaster*, open up a new opportunity of producing ‘first-in-the-class’ proteins towards functional endpoints. This study points to a large untapped genomic space from the fundamental biology and applications perspectives.

## Introduction

The genome is a fundamental hereditary chassis of an organism. It is a huge information repository that encodes molecular jobs in sequences to suit cellular requirements.

DNA sequences resemble information codes in a compressed format that get uncompressed at the time of transcription involving processes like alternative splicing, epigenetic modifications, and so on. The Encyclopedia of Coding Elements project (ENCODE) has helped us understand DNA sequences along the lines of functional assignments e.g., ‘made-for-RNA’, ‘made-for-proteins’, ‘no work assigned’, and ‘retired from employment’.

Among several interesting discoveries, one of the key findings is that a tiny portion of the genome is protein encoding, a large proportion of the genome is RNA encoding and the rest of the genome is naturally non-expressing (Feingold et al., 2004). The exact job assignment of sequences varies among organisms and is one of the key reasons for their immense functional diversity.

It is interesting that some sequences are expressed to their full potential and assigned protein coding jobs. A large proportion of sequences are partially expressed (only RNA encoding), and some are never expressed (intergenic sequences), or had a history of expression (pseudogenes). Sequences whose potential has been partially realized, never realized, or retired from expression jobs are collectively referred to as the Dark Genome (Dhar, 2022).

In 1972, Susumu Ohno, a famous Japanese Scientist introduced the term ‘junk’ in one of the prominent scientific meetings (Ohno, 1972). Nowadays, it is well established that what was previously considered as genomic debris is, in fact, RNA coding sequences (Saha et al., 2002).

Though protein-encoding and RNA-encoding sequences have received significant attention in the past, the ability of dark genome sequences to synthetically encode novel biomolecules has received little attention (Dhar et al., 2009, Shidhi et al., 2015, Raj et al, Chakrabarti et al., 2020).

To fill in the information gap, sequences in reverse orientation to the existing protein sequences were considered in this computational study. The reverse coding possibility was examined from the physiochemical property, structure, and function perspectives to build a use case for experimentally validating first-in-the-class molecules.

## Material and Methods

The Escherichia coli (Strain: K-12 MG1655) genome database and Saccharomyces cerevisiae (S288C) genome database and Drosophila melanogaster (fruit fly) genome database was considered in this study (NCBI Resource Coordinators, 2013). A total of 4315 protein-coding genes of E. coli, 6017 protein-coding genes of S. cerevisiae and 13,962 protein-coding genes of the Drosophila were retrieved from the NCBI database (https://www.ncbi.nlm.nih.gov) to generate reverse DNA sequence.

### Translation of reverse sequence

Sequences were computationally reversed and translated into proteins using the Unipro UGENE-Integrated Bioinformatics Tool (Okonechnikov et al., 2012). Full-length translated sequences without intervening stop codons were considered for further study. Sequences that showed stop codons were discarded and not taken for further analysis. The Expasy-Translation Tool (Gasteiger, 2003) was used to quality-check the result.

### Protein Sequence Similarity

The full-length reverse sequences were BLASTed against the NCBI Non-Redundant protein database for similarity assessment (Altschul et al., 1990). Only those sequences were considered for further study that did not show any resemblance with the known proteins (Table 2).

### Physicochemical properties

Using the Expasy Protparam tool of SIB Swiss Institute of Bioinformatics (Roy et al., 2011, physiochemical properties like instability index, aliphatic index, theoretical Pi, grand average of hydropathicity (GRAVY) index and molecular weight were computed (Table 4).

### mRNA secondary structure

mRNA secondary structure and free energy were calculated by using Mfold server (Zuker, 2003). Combining free energy from various secondary structural components yields an estimated total free energy of a molecule structure. The Mfold calculates this free energy and compares it with the relative thermodynamics of different structures from the structure of the same sequence.

### Structure, Function Prediction, Subcellular localization

The tertiary structure of proteins along with the probable function was predicted using the I-TASSER Server (Iterative Threading ASSEmbly Refinement) tool (Zhang, 2008). I-TASSER predicts with good accuracy and generates 3D protein structures. WoLF PSORT tool was used to compute subcellular localization (Horton et al., 2007). Ramachandran plot was used to calculate the spread of amino acids in the 2-dimensional space by calculating torsional angles of amino acids φ (phi) and ψ (psi) in a protein sequence. Stereochemical properties were studied using the Procheck server.

## Results

### Translation of Reverse Sequences

On reading protein sequences backward and translating all possible sequences in *E. coli, S*.*cerevisiae* and *D*.*melanogaster*, a small proportion of full-length reverse protein sequence equivalents showed up without stop codon (Table 1).

**Table 1.** Full-length reverse proteins in E. coli, S. cerevisiae & D. melanogaster.

| S. No. | Types of sequences | *E. coli* | *S. cerevisiae* | *D. melanogaster* |
| --- | --- | --- | --- | --- |
| 1 | Protein coding genes | 4315 | 6317 | 13962 |
| 2 | Full length reverse proteins | 46<br>(1%) | 31<br>(0.5%) | 103<br>(0.7%) |

### Sequence Similarity

The full-length reverse protein sequences from *E. coli, S. cerevisiae* and *D melanogaster* were matched against the existing NCBI protein sequence database. Table 2 summarizes sequence matching data of reverse proteins against all the non-redundant proteins from the NCBI database.

**Table 2.**
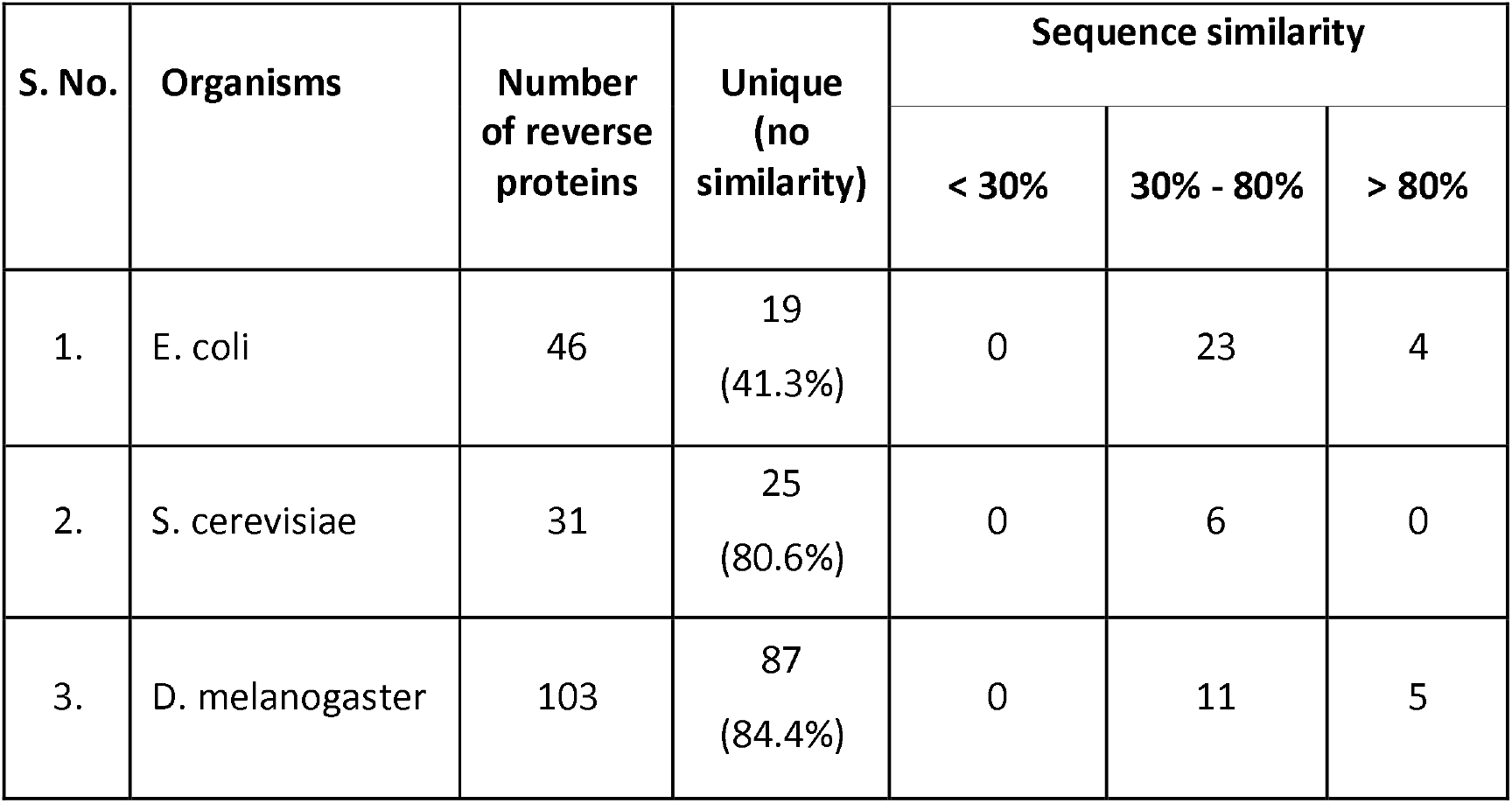
Sequence similarity data of reverse proteins from *E. coli, S. cerevisiae* and *D. melanogaster*.

Even though reverse protein sequences are completely different from the known protein sequences, results from Table 3 show high structural similarity with existing proteins in all three organisms studied (structural similarity score ranging from 50% to 90%). The I-TASSER values of tertiary structure prediction showed a favorable TM-score range i.e., above 0.5.

**Table 3.**
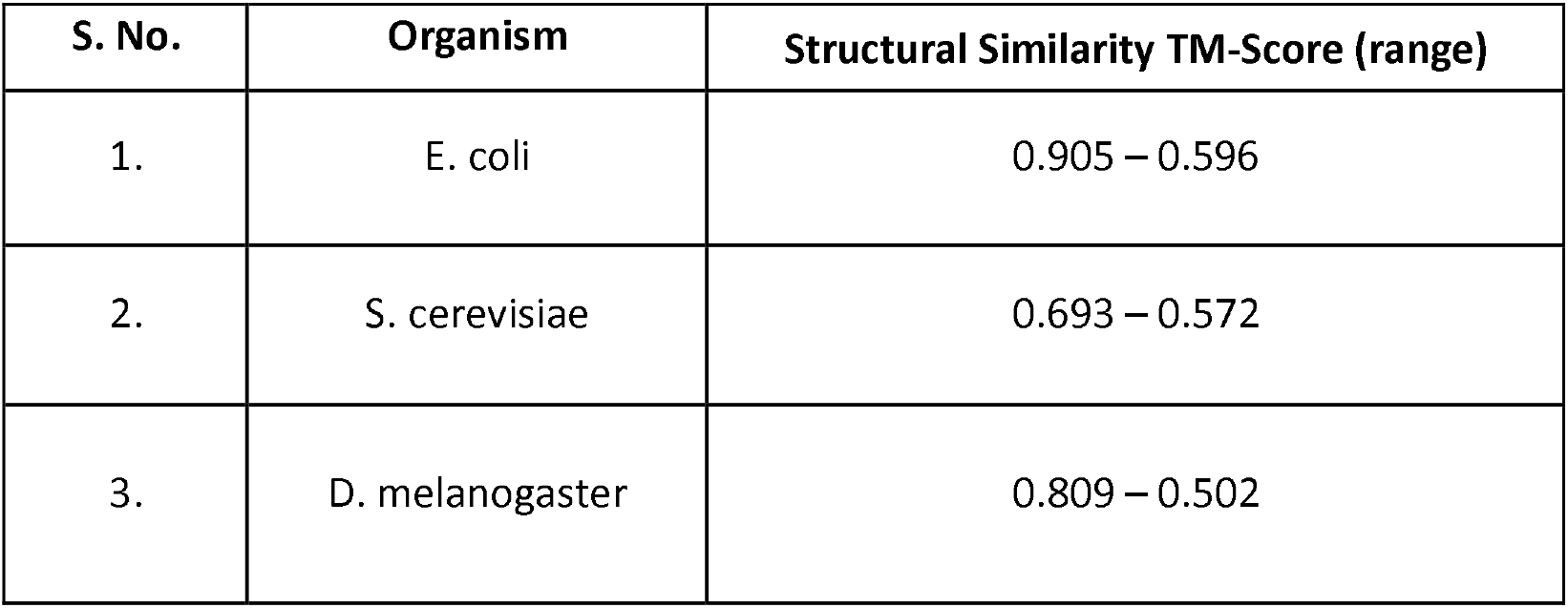
Predicted structural similarity of putative and unique reverse proteins of *E. coli, S. cerevisiae* and *D. melanogaster*.

### Physicochemical properties

A study on the physicochemical properties of reverse proteins (Table 4) indicates structural stability of proteins within a cell, if expressed. Most of the reverse proteins were found in the favorable range of isoelectric point, instability index and hydropathicity value. The codon usage defined by GC percentage was within the natural range found in three organisms.

**Table 4.** Physicochemical properties of putative reverse proteins of *E. coli, S. cerevisiae* and *D. melanogaster*.

| S. No. | Physiochemical property | *E. coli* | *S. cerevisiae* | *D. melanogaster* |
| --- | --- | --- | --- | --- |
| 1 | Length (amino acid) | 33 - 172 | 31 - 124 | 30 - 188 |
| 2 | Molecular weight (kDa) | 3.8 - 21.1 | 3.4 - 14.0 | 3.6 - 23.3 |
| 3 | Theoretical pI | 4.59 - 12.23 | 5.35 - 12.18 | 4.21 - 12.75 |
| 4 | Instability index | 11.26 - 91.22 | 11.88 - 94.57 | 7.86 - 217.7 |
| 5 | Grand average of hydropathicity (GRAVY) | -0.893 - 1.164 | -1.435 - 1.297 | -3.083 - 1.14 |
| 6 | G-C % (range) | 42.4 - 60.1 | 23.5 - 49.8 | 45.4 - 58.3 |

### mRNA secondary structure

The free energy of RNA secondary structure is determined by the combined contributions of its length, nucleotide content and nucleotide order. The one with the most negative value of free energy is considered the most structured and the most stable. All the reverse proteins were found to be in the favorable range (Table 5).

**Table 5.** mRNA secondary structure free energy of putative reverse proteins.

| S. No. | Organisms | mRNA secondary structure free energy (Kcal/mol) (range) |
| --- | --- | --- |
| 1 | E. coli | -1.664 to -23.50 |
| 2 | S. cerevisiae | -112.4 to -25.7 |
| 3 | D. melanogaster | -135.2 to -33.90 |

### Structure and Function Prediction

Many reverse proteins showed enzymatic function, and some were predicted to be DNA-binding proteins (Table 6). Normalized B-factor measurements through I-TASSER are predicted for the dynamic motion of proteins in the crystalline state. Most residues below 0 indicate the stability of proteins (fig 1).

**Table 6.** Predicted structure and function prediction of putative reverse proteins of E. coli, S. cerevisiae and D. melanogaster.

| S. No. | Organisms | Structure prediction (C-score) | Function prediction | Subcellular localization |
| --- | --- | --- | --- | --- |
| 1 | *E. coli* | -0.94 to -5 | lyase, oxidoreductase, transferase, electron transport, ATP synthase, aminotransferase | Integral membrane, cytoplasmic, nucleus, mitochondria, extracellular |
| 2 | *S. cerevisiae* | -1.3 to -4.3 | Transferase, hydrolase, lyase, oxidoreductase, Electron transport | Nucleus, mitochondria, vacuole, endoplasmic reticulum, plasma membrane-bound |
| 3 | *D. melanogaster* | -2.46 to -4.97 | Transport protein, oxidoreductase, transferase, membrane | Secreted, cytoplasmic, nuclear, extracellular |

**Fig. 1.**
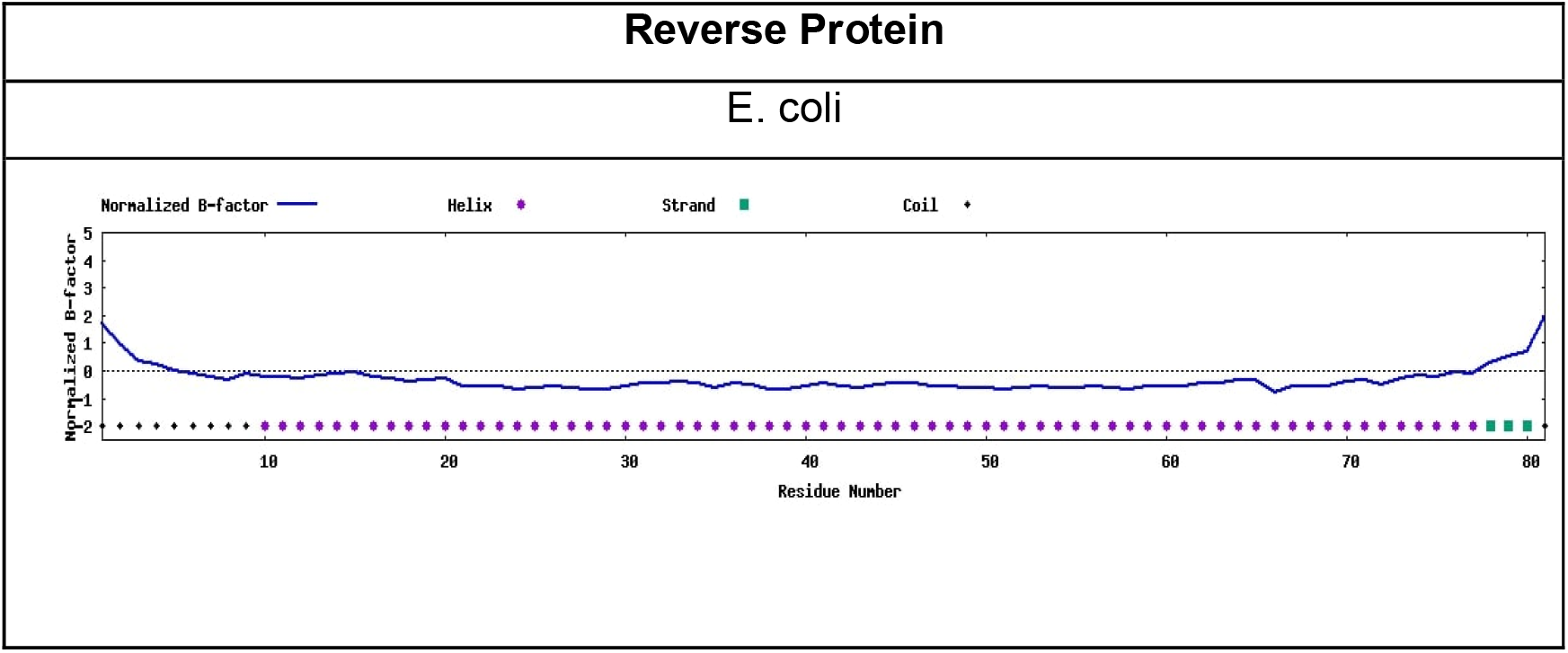
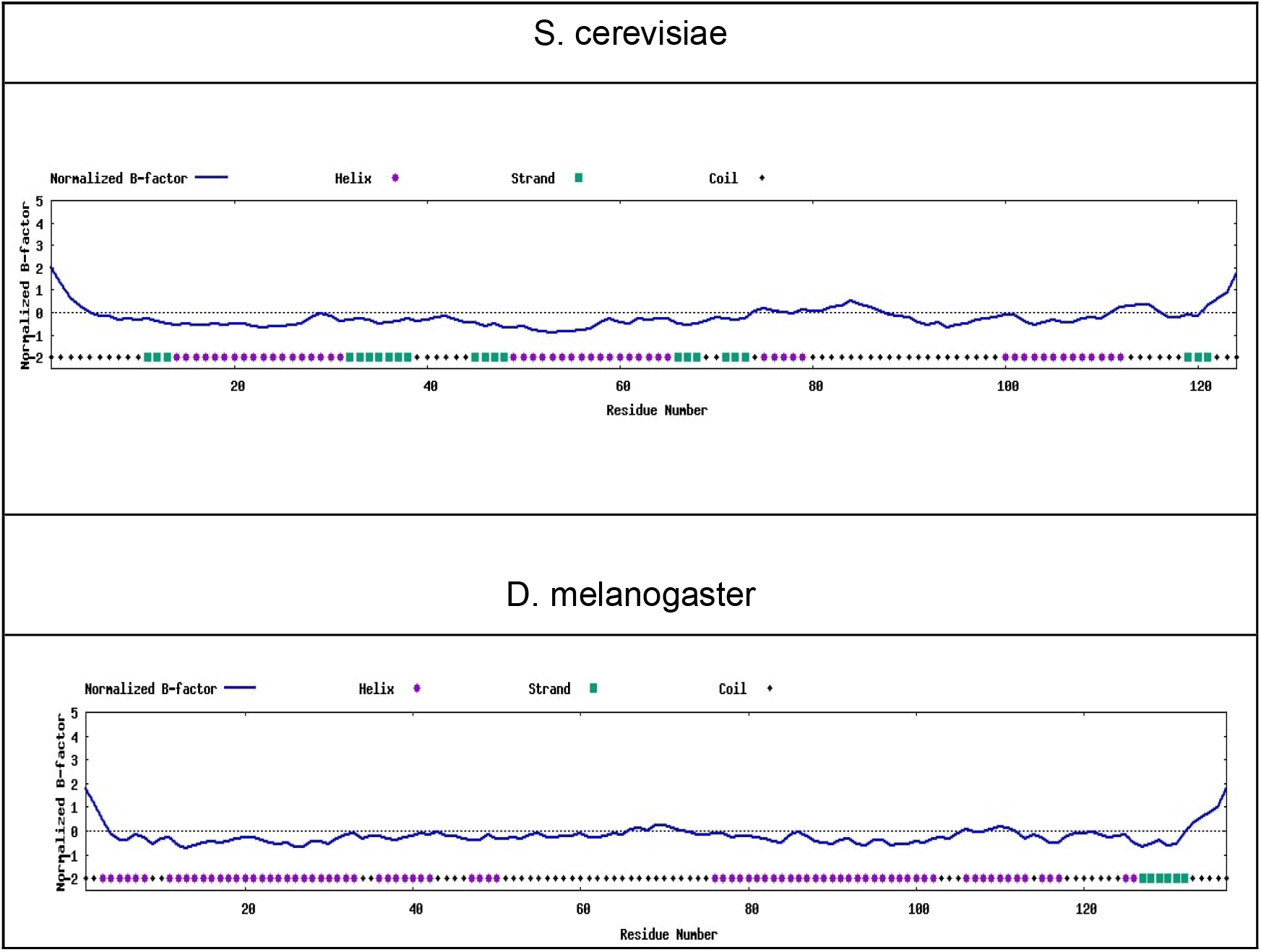
Normalized B-Factor Profile of reverse proteins in E. coli, S.cerevisiae and D.melanogaster.

**Fig. 2.**
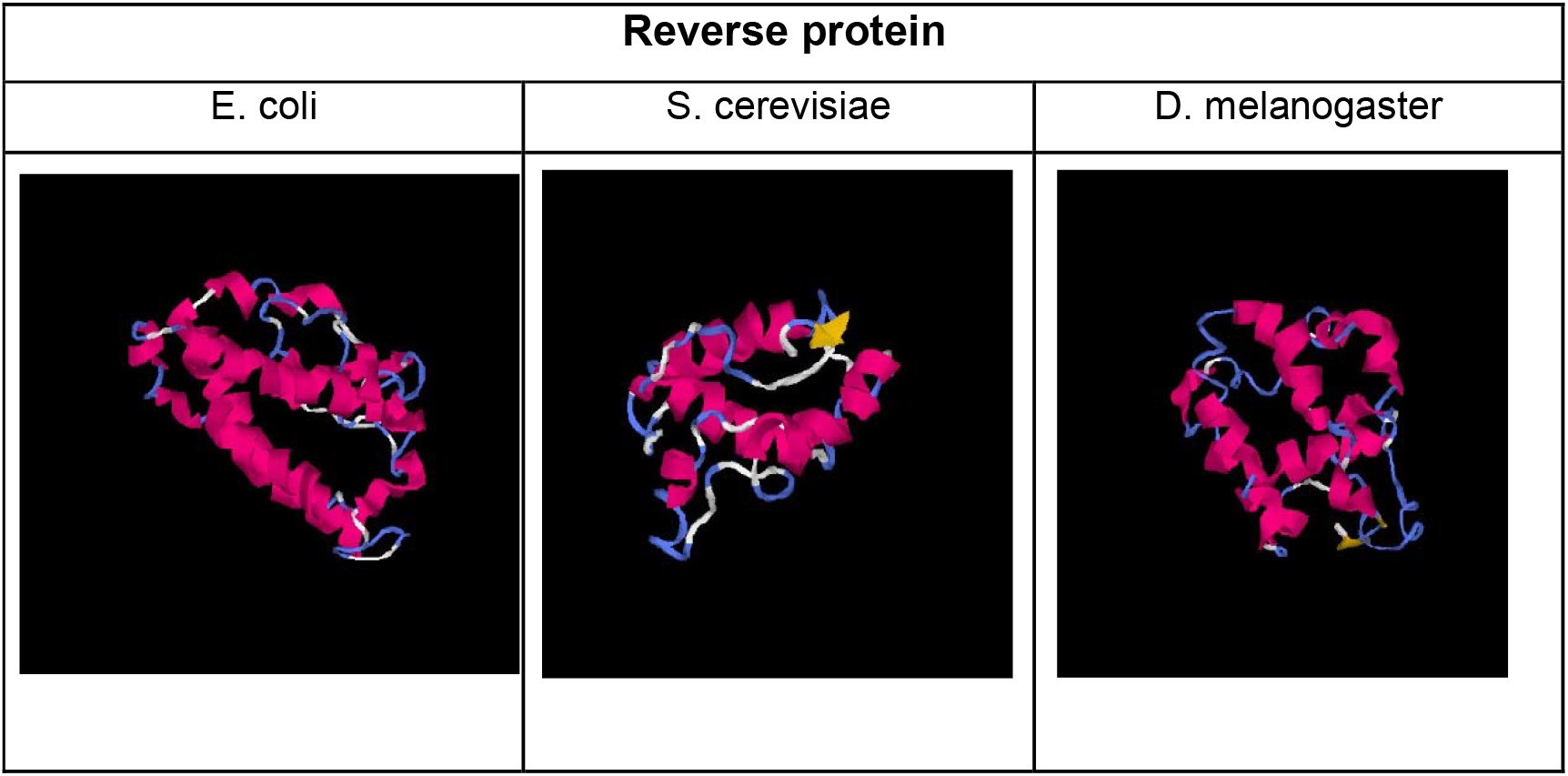
Tertiary structure of reverse protein sequences resembling naturally expressing proteins (prediction tool: I-TASSER)

The WoLF PSORT predicted protein localization. The location of most of the proteins were mapped to the cytoplasm, extracellular matrix, mitochondria, nucleus and membrane. Some of the reverse proteins were predicted to be secretory in function (Table 6).

### Stereochemical properties

The stereochemical property was studied using the Ramachandran plot (Fig 3). Most of the residues were predicted to be in the favorable regions indicating a good protein folding possibility.

**Fig. 3.**
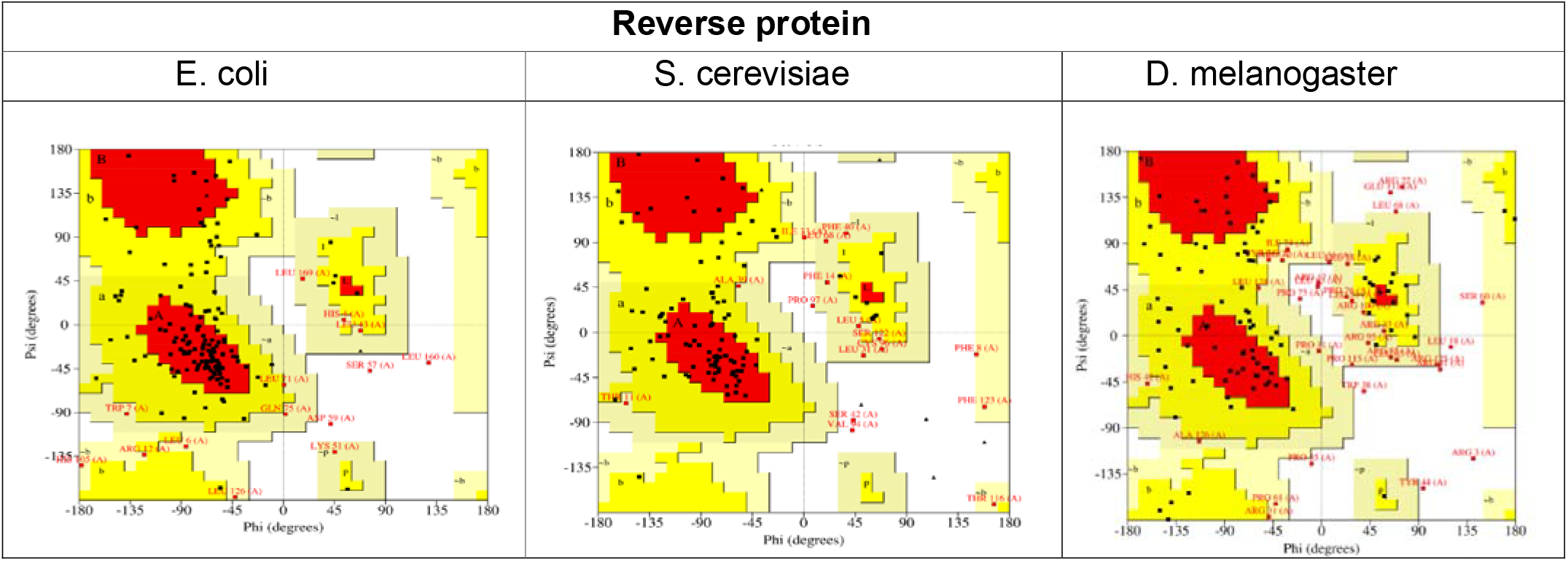
Ramachandran Plot of reverse protein sequences of E. coli, S. cerevisiae and D. melanogaster.

## 4 Discussion

The information encoded in a genome is a combination of an evolutionary hand-me-down format and horizontally installed plugins from other organisms. DNA sequences are assigned unique molecular jobs that allow the cell to run metabolic and regulatory pathways in a seamless and integrated manner. Of various types of genomic sequences, non-expressing DNA is the most unexplored region from structural, interaction, and functional perspectives.

The present reverse protein study derives its genesis from the dark genome paper that provided proof of the concept of making functional genes and proteins from intergenic sequences (Dhar et al., 2009). Subsequently, dark genome sequences were artificially expressed in *E*.*coli, D*.*melanogaster* and *S*.*cerevisiae* leading to antimalarial, antimicrobial and anti-cancer molecules (Joshi et al., 2013, Raj et al., 2015, Deepthi et al., 2017). Recently, tRNA sequences were used to produce tREPs (tRNA encoded peptides) showing a strong anti-leishmania properties (Chakrabarti et al., 2022). in this study, sequences of known protein-coding genes in *E*.*coli, S*.*cerevisiae* and *D*.*melanogaster* were computationally reversed leading to the identification of full-length reverse proteins. A new synthetic proteome inventory was created to include full-length protein sequences read backward mapping to less than 1 percent of known protein-coding genes in all three organisms. Computational structural studies on this small sample of reverse sequences indicated their stability at the mRNA and protein levels.

From the functional perspective, the non-expressing and partially expressing dark genome sequences led to novel enzyme predictions e.g., oxidoreductase, lyase, transferase, hydrolase and some ATP synthase. Most of the reverse proteins were predicted to have specific cellular addresses.

This study introduces the concept of reading sequences backward and generating molecules in the form of reverse proteins. Computational analysis of reverse proteins led to several interesting structural and functional predictions that merit experimental validation. Though the current study considers only the full-length reverse proteins, many possibilities would open if partial-length reverse proteins are also considered.

The current work provides the foundation to build a new synthetic proteome inventory with computational and experimental data. We are extending the study to include other organisms with the hope to understand evolutionary decisions and generate applications. In the natural setting, a bidirectional expression design seems to be a non-intuitive possibility. However, one may synthetically generate protein variants by reading sequences backward and adding more information to the genomic, transcriptomic, and proteomic inventories leading to applications.

## Author Contributions

PKD conceived the idea of making functional proteins from reverse protein-coding DNA strands. Further, he supervised the work and wrote the final version of the manuscript. SN performed all the computational work and drafted the first version of the manuscript.

## Acknowledgment

SN would like to express sincere thanks to UGC-CSIR for providing NET-JRF Scholarship. PKD warmly thanks JNU Administration for the critical lab support and Prof. Binay Panda (JNU) for very helpful discussions.

